# Decomposing metabolite set activity levels with PALS

**DOI:** 10.1101/2020.06.07.138974

**Authors:** Karen McLuskey, Joe Wandy, Isabel Vincent, Justin J.J. van der Hooft, Simon Rogers, Karl Burgess, Rónán Daly

**Affiliations:** Glasgow Polyomics, University of Glasgow, Glasgow, UK; IBioIC, Strathclyde Institute of Pharmacy and Biomedical Sciences, University of Strathclyde, Glasgow, UK; Bioinformatics Group, Department of Plant Sciences, Wageningen University, Wageningen, The Netherlands; School of Computing Science, University of Glasgow, Glasgow, UK; Centre for Synthetic and Systems Biology, School of Biological Sciences, University of Edinburgh, Edinburgh, UK

## Abstract

**Motivation:** Related metabolites can be grouped into metabolite sets in many ways. Examples of these include the grouping of metabolites through their participation in a series of chemical reactions (forming metabolic pathways); or based on fragmentation spectral similarities and shared chemical substructures. Understanding how such metabolite sets change across samples can be incredibly useful in the interpretation and understanding of complex metabolomics data. However many of the available tools suitable for the enrichment analysis of metabolite sets are based on simple methods that badly handle the missing features inherent in untargeted metabolomics measurements and can be difficult to integrate into existing applications.

**Results:** We present PALS (Pathway Activity Level Scoring), a Python library, command-line tool and Web application that performs the ranking of significantly-changing metabolite sets over different experimental conditions. As example applications, PALS is used to analyse metabolites grouped as pathways and by common MS-MS fragmentation structures. A comparison of PALS with two other commonly used methods (ORA and GSEA) is also given, and reveals that PALS is more robust to missing peaks and noisy data than the alternatives. We report results from using PALS to analyse pathways from a study of Human African Trypanosomiasis. Finally, we also report how PALS used tandem MS fragmentation structures to reveal enriched metabolite sets between clades in Rhamnaceae plant data, and on American Gut Project data.

**Availability:** PALS is freely available from our project Web site at https://pals.glasgowcompbio.org/. It can be imported as a Python library, run as a stand-alone tool or used as a web application.

## 1 Introduction

An organism’s metabolism describes all of the chemical reactions involved in its cells, where the small-molecule intermediates and products of these reactions are known as metabolites. Untargeted metabolomics provides a profile of all detectable metabolites in a system, allowing the unique chemical fingerprint left behind by the metabolism to be investigated. Such studies are imperative in understanding changes in biological mechanisms related to environmental and genetic variations. Mass-spectrometry (MS) is one of the most commonly used techniques for untargeted metabolomics and is often coupled with chromatographic separation using liquid chromatography (LC). Experimental data from a typical LC-MS experimental sample is a series of spectra, where signal intensities are generated for each detected ion. After data pre-processing, a table of chromatographic peaks is produced, where each peak can be represented by its m/z, RT and intensity values. Identification is performed to associate peaks with metabolite identities, either by matching m/z and RT with internal standards or through comparisons against mass spectral databases.

Understanding how groups of metabolites are affected by experimental factors can be informative to gaining biological insights. Rather than analysing individual metabolites in isolation, often sets of related metabolites are considered. One way to define metabolite sets is through prior knowledge of how metabolites participate in a series of related chemical reactions (comprising metabolic pathways). Ranking biologically-relevant pathways through changes of associated metabolite intensities provide high level information that helps prioritise relevant pathways [16]. An alternative way to group metabolites is to utilise the wealth of structural information present in fragmentation spectra from tandem mass spectrometry. Molecular networking [31] uses spectral similarities to group metabolites into *Molecular Families* (MF). These MFs potentially correspond to chemical classes and have proven invaluable in enhancing the putative identifications of unknown metabolites [9, 14]. In MS2LDA, metabolites are grouped by co-occurring fragments and neutral losses into *Mass2Motifs* that reveal potential chemical substructures [28].

In recent years, enrichment analysis algorithms, originally developed for analysing large collections of genes or transcripts, have been adapted to metabolomics. These methods were traditionally used to test gene expression data for functions or processes that are over-represented (i.e. enriched) with regard to functional gene sets, pathways and networks. Considering differentially expressed (DE) gene sets, two of the most commonly used types of gene enrichment techniques are: Over-Representation Analysis (ORA) and Functional Class Scoring (FCS). ORA approaches rely on testing for the presence or absence of significantly changing genes in a pathway, using statistical significance tests. FCS methods, which include Gene Set Enrichment Analysis (GSEA) [25] use statistical approaches to identify gene sets that are significantly enriched or depleted in the pathway. FCS improves upon ORA by eliminating the need to preselect significantly changing genes, as well as taking into account how groups of genes can be co-expressed together (instead of assuming they are independent as in ORA).

In metabolomics, a variety of methods that encompass enrichment analysis of DE metabolite sets (in particular pathways) have been developed. These includes comprehensive platforms such as MetaboAnalyst [4] and MeltDB [15], both of which use an implementation of Metabolite Set Enrichment Analysis [23], a variant of GSEA. At the time of writing, MetaboAnalyst is arguably the most popular online tool for pathway ranking [4]. Various stand-alone tools are also available with many of them implemented in the R programming language, e.g. MetaboAnalystR [3], or available as Web-based systems, e.g. ImPaLA [12].

Benchmarking studies comparing these tools are relatively rare. This is in part a result of the paucity of metabolomic studies where the ground truth of significantly changing pathways is known. However, it is possible to look at studies comparing gene set enrichment analysis on microarray expression data. One such comprehensive study compares gene set analysis methods using prioritisation, sensitivity and specificity [26]. According to this study, the best ranking method with low false positive rates is Pathway Level Analysis of Gene Expression (PLAGE) [27]. PLAGE performs singular value decomposition (SVD) to compute a pathway activity value from expression data in a sample. This has advantages over other methods in that the PLAGE method is computationally simple while returning high performance in both sensitivity and specificity. Decomposing activity levels via SVD generalises well and appears particularly relevant when applied to MS peak intensity values. This was demonstrated in our previous work [28] where PLAGE was successfully used to assess DE Mass2Motifs in different beer samples. However, we note that in general PLAGE has hardly been used by the metabolomics community to assess the activity level of metabolite sets and pathways.

This manuscript introduces **PALS** (**P**athway **A**ctivity **L**evel **S**coring), a tool for ranking of pathways centered around peak intensities and compound annotations based on the matrix decomposition method in PLAGE. PALS is designed to look at how metabolic pathways or metabolite groups change between different experimental conditions by decomposing peak intensities in a pathway into latent factors corresponding to observed metabolic changes (activities). It rates small correlated changes in a group of pathway metabolites as more interesting than large changes in an individual metabolite, where other metabolites in the pathway are not changing. The changes can be increases and/or decreases in the peak intensities, but must occur as part of the pathway group. This is particularly important in situations where enzyme inhibition results in substantial increases to metabolite abundance before a lesion in the pathway, with concomitant decreases after the lesion (see [29] for an example). The modularity of PALS also means it is not limited to pathways, and any user-defined grouping of metabolites into metabolite sets can be readily analysed. We demonstrate this by assessing the activity levels of metabolite sets grouped by similarity of fragmentation spectra.

## 2 Methods

### 2.1 Running PALS

PALS can be run either as a stand-alone tool or imported as a Python library, allowing users with a wide range of expertise and requirements to use it (input format detailed in Supplementary Section S1). For beginners, a Web interface is provided to run PALS in a user-friendly manner (refer to Section 2.4 for more details). For intermediate users, PALS can be run as a stand-alone command-line tool and be incorporated into a workflow as part of a custom script. In this mode, peak and annotation CSV files are input on the command line along with experimental design parameters, and a variety of other available parameters, accessible through command-line options (Supplementary Section S2). Pathway ranking results are output as a CSV file-listing pathways and their p-values. Expert users can also incorporate the PALS library directly in their Python application or use it in an interactive data analysis environment such as Jupyter Notebook [18]. In programmatic usage, the required input can be passed as data structures directly to the PALS object. A comprehensive tutorial describing how to use PALS as a library is available from the project Web site. Along with being available through a Python library and Web application, PALS is also integrated into several projects such as PiMP [10], FlyMet (flymet.org) and WebOmics (webomics.glasgowcompbio.org).

### 2.2 Pathway Database Queries

The presence of isomers and isobaric compounds in a sample means that multiple peaks with close m/z values can be annotated with the same molecular formula although those peaks are actually produced by different compounds in the MS. Resolving the actual molecular identities of annotated peaks is a challenging problem even at very high resolution [17]. To ensure coverage, PALS represents compounds using their chemical formulae when mapping peaks to pathways. As such, all peaks having the same formula annotations are used for activity level analysis of a pathway (Figure 1A).

**Figure 1:**
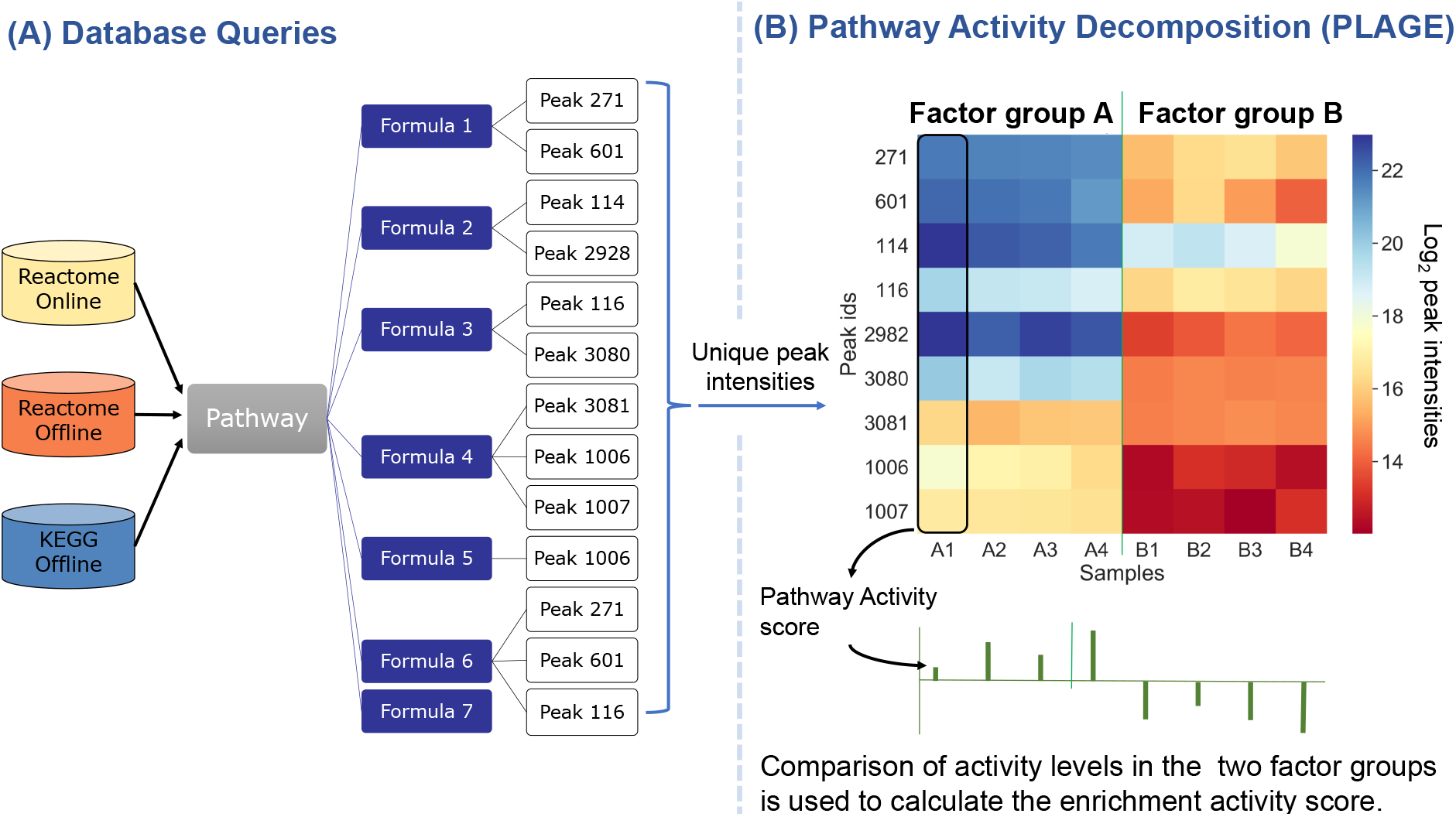
Overall schematic of PALS. **(A)** Different databases (Reactome Online, Reactome Offline and KEGG Offline) can be used to query pathways. Peaks are mapped to pathways through their formula annotations. For a set of compounds in a pathway, all peaks sharing the same formula annotations of those compounds are used to construct the intensity matrix. **(B)** In PLAGE, the intensity matrix is decomposed via SVD to obtain the enrichment activity score of the different factor groups for that pathway.

When PALS runs in offline mode, metabolites, pathways and the mapping of metabolites to pathways are collected from an existing dump of the KEGG database [13] used in PiMP [10] or from the metabolic pathways in Reactome [6] exported for 18 common species. These dumps store the information on compounds and their pathway mapping for the given species as a compressed JSON format. To utilize the most recent version of Reactome, PALS can also be run in online mode that queries pathways directly from a Reactome instance running on top of a local Neo4j graph database. Queries are created in PALS using Cypher (the graph query language used for Neo4j) to retrieve the latest pathways and the mapping of compounds to pathways from the database. Reactome is regularly updated, with the entire database available from the Reactome website.

### 2.3 Decomposing Activity Levels

The initial activity level score for a pathway is computed using singular value decomposition (SVD) following PLAGE [27], and is briefly summarised here. Similar to the way that gene expression data can be decomposed into meta-genes and activity levels in PLAGE, the intensity matrix *X* associated with a pathway can also be decomposed via SVD to obtain that pathway’s activity score (Figure 1B). Given an intensity matrix *X* (where rows are the peaks annotated with formulae found in the pathway, and columns are the samples in the dataset), the decomposition of *X* can be written as *X* = *U*Σ*V* where columns in *U* are the orthonormal left singular vectors representing ‘meta-compounds’, Σ contains the diagonal matrix of singular values arranged in descending order, and rows in *V* are the right singular vectors representing ‘meta-samples’. Enrichment activity (EA) score in a sample is given by the first row *v* in *V*, which is a vector of coefficients representing the activity levels of the first meta-compound in *U* (having the largest singular value) across samples. For more details, see [27].

To perform pathway analysis, the EA scores obtained from the PLAGE procedure are used to calculate the t-statistic for each pathway for the two experimental factors being compared. As in [27], random permutations of the sample labels are performed to obtain the null distributions of *t*-statistics. Here an enhancement to the original PLAGE approach is introduced in order to obtain better calibrated enrichment activity results: during the permutation test, it is observed that the tail end of the distribution of permuted *t*-statistics occasionally contains some extreme values that could skew results. To improve the stability of these results, the minimum and maximum *t*-statistics from each permutation test are modelled using the Generalised Extreme Value (GEV) distribution [1], with parameter fitting using maximum-likelihood estimation. The *t*-statistic of a pathway is then evaluated against the fitted density in order to produce the final EA score of a pathway.

### 2.4 Interpretation of Results

To assist in results interpretation, *PALS Viewer* provides a user-friendly Web-based interface to run pathway analysis and inspect the results (see Supplementary Section S2.2). It can be run locally or accessed from the project Web site. From the sidebar, users upload their intensity and annotation CSV files and configure user-defined parameters including experimental design, preferred species and database. Pathway decomposition is run, and analysis results are shown in an interactive pathway ranking table. Each entry in the table shows pathways, their corresponding PALS p-values and the number of formula hits. These can be used for sorting and filtering by chosen thresholds. Selecting a pathway from the Pathway Browser reveals the Reactome (or KEGG) pathway diagram. Information on the fold changes of annotated compounds are submitted to Reactome Analysis Service and mapped onto the pathway diagram for Reactome. As an option, users can also display a heatmap of the annotated formula hits in the pathway. Choosing to order the pathways by the p-values means small but consistent changes in a groups of pathway metabolites should appear near the top (most interesting), even if the changes in the individual metabolites are modest.

### 2.5 Analysis of User-defined Metabolite Sets

The decomposition approach employed by PALS is not limited to the analysis of pathways. In fact any user-defined grouping of peaks where each metabolite set can be represented as an intensity matrix (peaks-vs-samples) can be used for activity level decomposition. We demonstrate this in PALS Viewer by letting users run two extra methods to define the grouping of their peaks in a data-dependent fashion (in contrast to pathways which rely on prior knowledge). The first method uses molecular networking [FBMN, 22] to cluster peak fragmentation spectra by their similarities and produce the groups of peaks known as Molecular Families (MF). The second method uses a topic-modelling approach [MS2LDA, 28] to generate groups of peaks (called Mass2Motifs) based on the shared presence of fragment and neutral loss features in their fragmentation spectra. Both FBMN and MS2LDA are accessible as workflows from Global Natural Products Social Molecular Networking [GNPS, 31], which provides community resources to run large-scale molecular networking and annotations of spectra.

To analyse MFs in PALS, users provide a link to an existing FBMN result containing the MS1 peak table (for the intensity matrix) and MF information (for the peak grouping). For Mass2Motifs analysis, users provide links to both an existing FBMN result (for the MS1 information) as well as to an MS2LDA result describing the grouping of peaks into Mass2Motifs. Data is loaded in PALS, and peaks are allocated to metabolite sets according to their groups (note that a peak can be assigned to only one MF but to multiple Mass2Motifs). Activity level decomposition is performed on the metabolite sets following Section 2.3. The results are presented in PALS Viewer in a similar manner to pathways (described in Section 2.4): MFs or Mass2Motifs are shown in a ranked interactive table next to their p-values. Upon selecting a metabolite set (either an MF or a Mass2Motif), a heatmap is displayed showing the intensity values of member peaks across samples, as well as any additional metadata retrieved from GNPS or MS2LDA.

## 3 Results

### 3.1 Synthetic Data Experiments

#### 3.1.1 Synthetic Data Setup

To assess the robustness of PALS, benchmarking experiments were initially performed using synthetic data. Seven pathways were constructed with significant changes (showing clear block structures between the case and control groups in their peak intensity matrix) between two experimental conditions, each consisting of four samples. Using a normal distribution, random noise was generated (Supplementary Section S3) and standardised for input (Supplementary Section S1). Each of the seven pathways contained a different number of formulae listed in the following order: {2,4,6,10,20,40,80} (a one-to-one correspondence between an observed peak and a metabolite is assumed). Synthetic pathways were labelled by the number of formulae assigned to them (e.g. pathway *twenty* has 20 formulae and therefore 20 peaks). In addition, 100 background pathways containing only noise (showing no significant changes between the case and control groups) were generated. To simulate missing peaks, which often occurs in real data, peaks were also randomly removed from pathways with a uniform probability of 0.2. The performance of PALS (where decomposition is performed using the PLAGE method) was compared to widely-used over-representation (ORA) analysis and Gene-Set Enrichment Analysis (GSEA) methods, described further in Supplementary Section S4. These alternative methods were used for benchmarking experiments and are included in the PALS Python library, allowing all methods to be compared by utilising the same database query codes and retrieving identical pathways from KEGG and Reactome.

#### 3.1.2 Evaluation

The following evaluation metrics were implemented to assess the performance of the different methods. Let *T*={*two, four, six, ten, twenty, forty, eighty*} be the set of true answers (the labels of seven pathways with the corresponding number of peaks that are actually changing). Different pathway analysis method *m* will return different *P_m_*, a ranking of significantly changing pathways under some threshold *t* (set to 0.05 following convention), where pathways with p-values below *t* are considered significant (positives), and otherwise. For a method *m* the following counts can now be evaluated: **True Positives** (TP) = in *P_m_* and in T; **False Positives** (FP) = in *P_m_* but not in T; **False Negatives** (FN): in *T* but not in *P_m_*. To obtain single-number summaries, precision and recall is used. Here precision (*Prec* = *TP*/ (*TP* + *FP*)) measures pathway ranking relevancy and it often occurs as a trade-off to recall (*Rec* = *TP*/(*TP* + *FN*)), which measures how many actually relevant changing pathways are returned. High precision suggests that a method has a low false positive rate, while a high recall suggests that the method has a low false negative rate. To summarise overall performance, the Fj score, which is a harmonic mean of precision and recall and defined as *F*_1_ = (2 * *Prec* * *Rec*) / (*Prec* + *Rec*), is used.

#### 3.1.3 Synthetic Data Results

In this experiment, an increasing number of noisy peaks, containing random intensity values, were added to the seven pathways comprised of significantly changing peaks. This scenario reflects the case of attempting to find pathways with high activity levels when the number of significantly changing metabolites are small compared to noisy data or non-changing metabolites. Noise levels are defined in an increasing order of severity from 0%, 25%, 50%, 100%, 250%, 500% and 1000% of the original number of peaks in the pathway. For example, adding 50% noisy peaks to pathway *forty* results in 40 significantly changing peak intensities + 20 non-changing peak intensities in the pathway. The resulting synthetic data matrix with the addition of noise is then used as input to the methods being compared. This procedure is repeated 500 times for each noise level.

Assessing changes to the p-values of the true changing pathways as noise levels are added reveals that increasing noise levels generally produces higher p-values (Figure 2). This suggests that it becomes harder for pathways to be identified as significantly changing when a greater number of noisy peaks are present. At noise level 0%, both ORA and PALS perform well returning median p-values of 2E-6 and 0.0049, respectively. GSEA performs less well with median p-value of 0.0578, which is close to the selected significant threshold of 0.05 (shown as dotted line in Figure 2). The mean p-values from all methods increase as the noise level is increased. At 100% noise level, both ORA and PALS perform well even when there are as many noisy peaks are there are actually changing peaks, returning median p-values of 0.0083 and 0.0003 for PALS and ORA, respectively. At higher noise level (250%, 500% and 1000%), GSEA appears most sensitive to noise, whilst PALS perform best amongst the methods compared.

**Figure 2:**
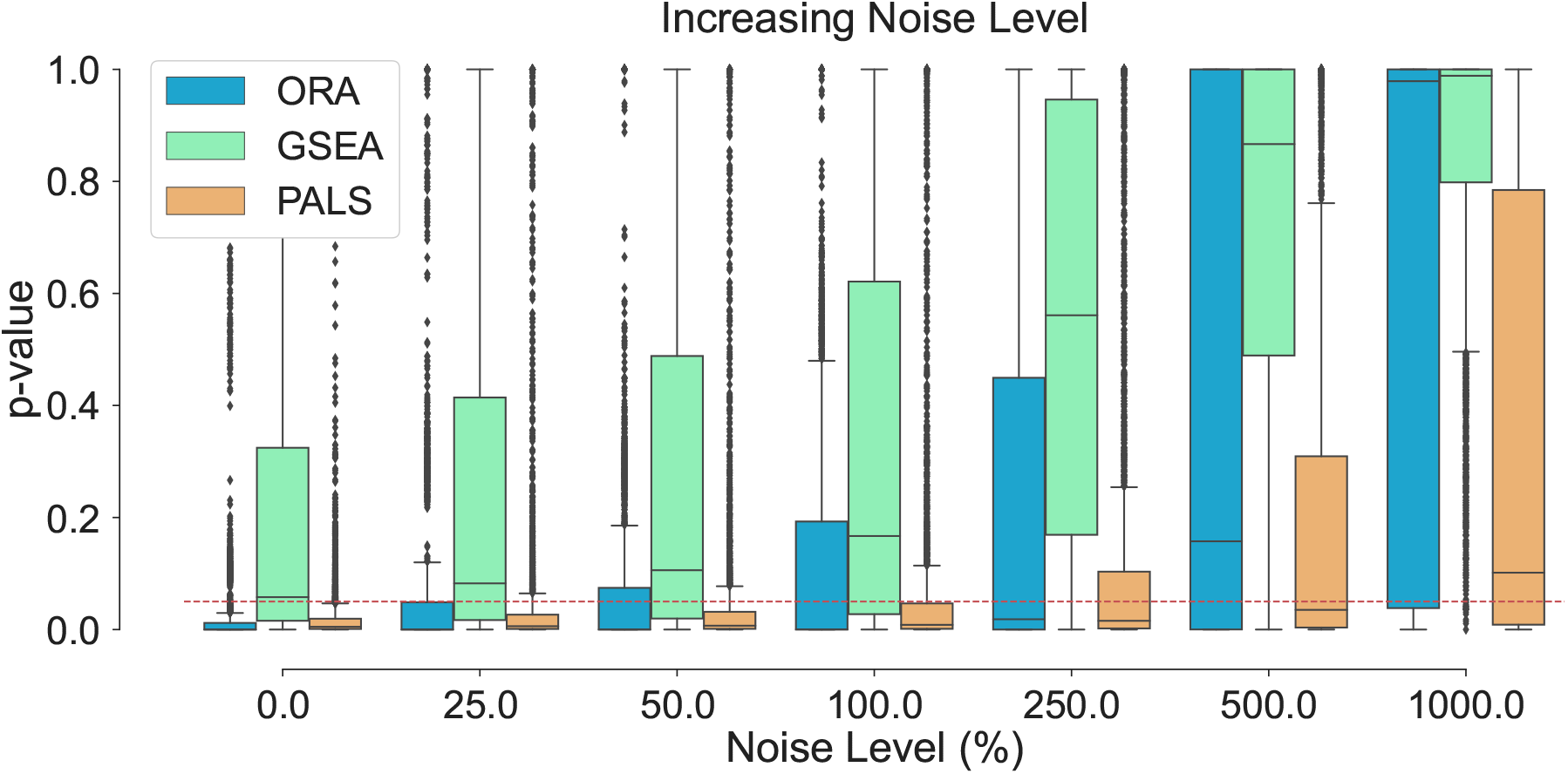
Adding random noise to the seven known changing pathways *T* = {2, 4, 6, 10, 20, 40, 80}. Noise level represents the number of noisy peaks added to a pathway as a percentage of the original number of peaks in that pathway. The boxplots show the spread of the p-values calculated from the pathways, including the median (solid horizontal line). The red dashed line indicates a p-value threshold of 0.05. The results show that increasing noise levels generally produces higher p-values making it harder to detect significantly changing pathways. PALS is more robust compared to other methods in returning lower p-values in the presence of noise.

Inspection of individual significant pathways of varying sizes (Figure 3) revealed that the mean p-values returned by all methods generally decreases with an increasing number of changing peaks in a pathway. This suggests that the greater the number of metabolites that are changing together, the higher ranked that pathway would be. As a result of the small number of metabolites in the pathway, all methods struggle to correctly identify pathways *two* and *four* as significantly changing at any noise level and generally perform better on the larger pathways *twenty, forty* and *eighty*. This reveals that the larger the number of changing peaks identified in a pathway, the more tolerant the method is to non-changing (random) peaks. For all pathways across all noise levels, PALS returns lower p-values compared to ORA and GSEA, which means more significantly changing metabolite sets can be detected.

**Figure 3:**
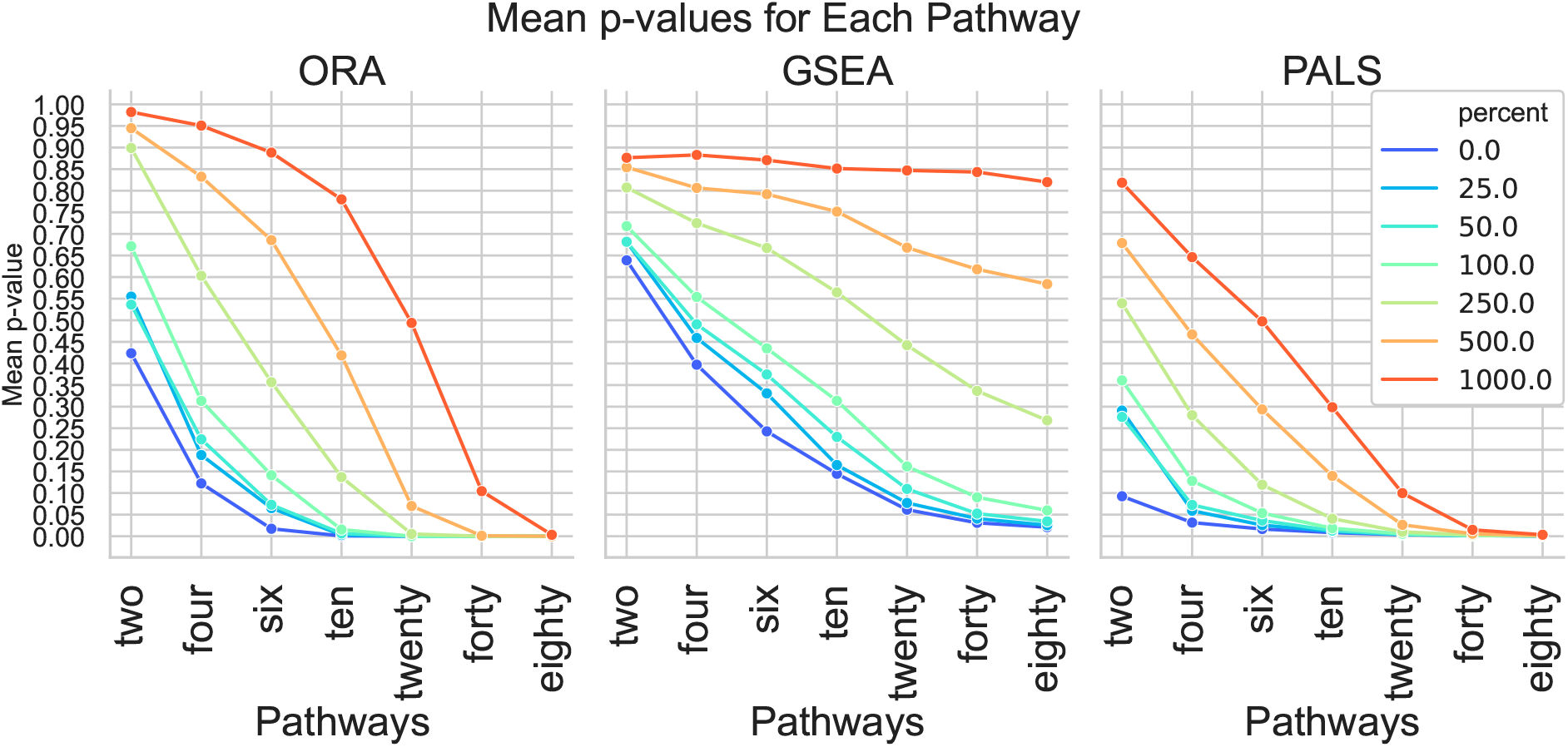
Mean p-values from ORA, GSEA and PALS calculated for each significantly changing synthetic pathway at different noise levels. Across all noise levels, it is easier to identify larger pathways as significantly changing than smaller pathways. PALS generally returns lower p-values than ORA and GSEA.

Finally, *F*_1_ score performance was evaluated. This summarises overall retrieval ability taking into account the number of true positives, false positives and false negatives of the methods being tested. Using this metric, it could be seen that the performance of ORA and PALS were roughly similar using 0-100% noise levels (Figure 4). At higher noise levels of 250%, 500% and 1000%, the *F*_1_ scores of PALS were consistently higher than ORA and GSEA. These results showed that a greater number of true positive pathways could be identified with PALS than with ORA or GSEA. GSEA performed worst among the three methods tested particularly at high noise levels, showing its sensitivity to noise. Additionally in Supplementary Section S5 we explored the effect of introducing an increasing number of missing peaks from the pathways, and broadly similar results are obtained with PALS outperforming the benchmark methods.

**Figure 4:**
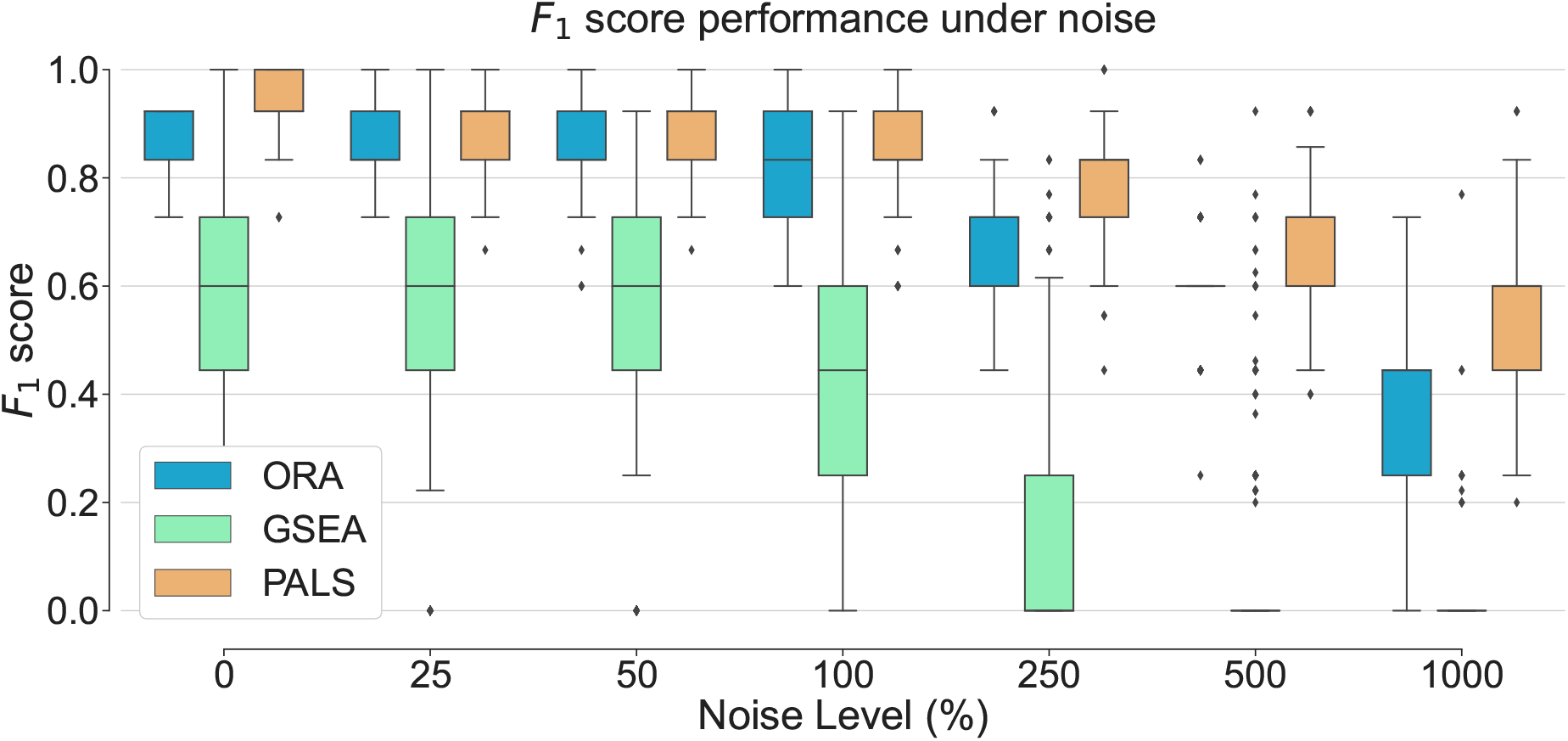
Overall performance of the different pathway ranking methods. Distribution of the *F*_1_ for each method under increasing noise conditions is shown with outliers displayed as small diamonds. The performance of PALS and ORA are roughly similar using 0-100% noise levels, but PALS performs better returning greater *F*_1_-scores at higher noise levels.

### 3.2 Real Data Experiments

To evaluate the performance of the proposed method on actual complex biological data, PALS was used to analyse metabolomics data obtained for a study on Human African Trypanosomiasis (HAT) introduced in [30]. The causative agent of HAT is the parasite *Trypanosoma brucei*, which is transmitted to a human/mammalian host by the bite of the tsetse fly. Two datasets of samples collected from human blood plasma and cerebrospinal fluid (CSF) are available from the study, denoted as the *Plasma* and *CSF* datasets. The control group consists of parasite-free patients, while the two case groups are those with stage 1 (S1) (parasites present in blood/lymphatics) or stage 2 (S2, parasites found in the CSF) trypanosomiasis.

#### 3.2.1 Case Study

Using the HAT data with PALS allows for the analysis of complex clinical data to be explored and demonstrates how pathway decomposition via PLAGE can be used to draw relevant biological conclusions. The Plasma and CSF datasets from the HAT study, comprising of 20/17 control (C) samples and 20 samples from both S1 and S2, respectively were uploaded and processed through the PiMP pipeline [10]. PALS was easily integrated into the PiMP pipeline and set to run automatically at the end of a data analysis workflow. PALS results from the HAT data revealed that a greater number of significantly changing pathways (p-value ļ 0.05) were found between the disease stages for the CSF than Plasma data (52/74; 42/128; 42/143 for Plasma/CSF when comparing S1:C; S2:C and S1:S2, respectively). In all of the comparisons, CSF samples have a greater number of changing pathways than those seen in the Plasma data. In particular, there is a notable difference in the changing pathways between the S1:C and S2:C/S1:S2 in the CSF. This can be explained by the fact that the parasites are only present in the CSF in S2 parasitemia and consequently more metabolic changes are expected in the CSF from S2 patients.

Looking at metabolic differences between S1 and S2 samples, the top-ranking PALS pathways in the CSF were heavily biased towards amino-acid metabolism (Supplementary Table S6). Changes in these pathways make sense for two reasons: Firstly, because all parasites scavenge nutrients from their hosts and *T. brucei* is predicted to be auxotrophic for the amino acids arginine, glycine, histidine, isoleucine, leucine, lysine, phenylalanine, tryptophan, tyrosine and valine [2], and secondly because there are unlikely to be parasites in the CSF of patients with a S1 infection. To support this finding, individual amino acids identified in the experiment were examined and it was revealed that all of the amino-acids required by *T. brucei* (listed above) (excluding glycine which was undetected) were found to be decreasing in the CSF of patients with S2 of the disease (Supplementary Table S7). Interestingly, running PALS on the Plasma data did not show any bias towards amino-acid metabolism, suggesting that the parasites potentially have a much more marked effect on the amino-acid biosynthesis and availability on their host when it reaches the CSF.

It is known that when *T. brucei* reside in the human bloodstream that they are exposed to high levels of glucose, which they rely on for their energy metabolism [5]. This reliance on glucose changes when the parasites are resident in the insect midgut, as they have a less readily available supply of glucose, and instead switch to amino-acids and their main carbon source [20]. Little is known about the metabolism of the parasites in the CSF, so it may be possible that amino acid usage is required to supplement the lower levels of glucose in the CSF (4.5 mM and 3mM in Plasma and CSF, respectively [32]). Another reason that uptake of amino acids is evident in the CSF samples but not in the Plasma samples could be a result of amino acid concentrations being much lower in the CSF (4-50 fold lower for the 9 amino acids than *T. brucei* are auxotrophic for [32]) and consequent salvaging of the nutrients by the parasites is more noticeable in the CSF.

#### 3.2.2 Real Data Experimental Setup

The HAT case study results demonstrate how PALS identified significantly changing pathways that can be explained and interpreted as biologically relevant. Here we assess the robustness of the different pathway ranking methods on real data. In a manner similar to the synthetic experiment, a range of missing peaks proportions *p* was set from 0.2 to 0.8, in increments of 0.2. After processing in the PiMP pipeline, the complete Plasma and CSF data produce 15584 and 8154 peaks, of which 1647 and 1132 have associated metabolite annotations, respectively. Therefore, *p* times the total number of peaks were randomly removed, and the rest used as input for pathway analysis. Assuming that a particular pathway ranking *P_m_* returned by a method *m* on the complete data will be better than the ones obtained from the reduced data (with missing peaks), results from the complete data could be used as the true answers for evaluation. Here true answers *T* are restricted to be the set of significant pathways above a threshold of 0.05 for their p-values of PALS and ORA, and a larger threshold of 0.25 for GSEA (following the suggestion in [25]). Given *T* and *P_m_*, performance in terms of precision, recall and F1 score can be computed as described in Section 3.1.2. This evaluation procedure was repeated 500 times. Comparisons were performed between the Stage1/Control groups on the Plasma data and between the Stage2/Control groups on the CSF data where more changing pathways are expected to be found in the dataset between the case and control groups as the disease progresses. The results are reported in Section 3.2.3.

#### 3.2.3 Robustness on Real Data

For all methods, it can be noted that the *F*_1_ score decreases as the proportion of missing peaks increase (Figure 5). This suggests that it becomes harder for all tested methods to reconstruct the original pathway ranking results obtained from the full data when fewer input peaks are available. On both the Plasma and CSF data, pathway decomposition in PALS performs best demonstrating the most robust tolerance to missing peaks, followed by ORA. In both datasets, GSEA performs worst as a result of its sensitivity to noise. The results obtained for GSEA on the HAT data agrees with the synthetic data experiment.

**Figure 5:**
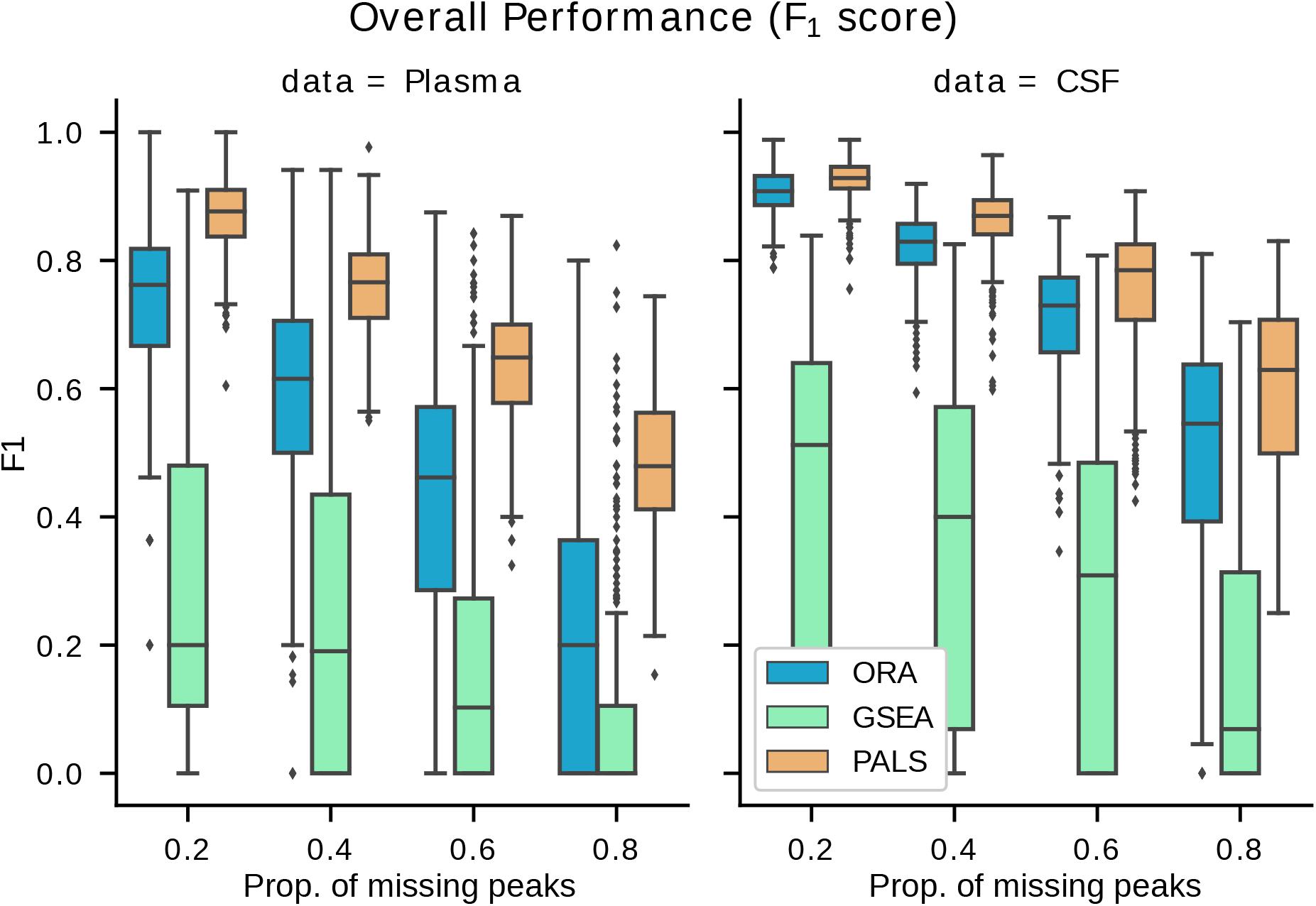
F1 score results from the HAT data experiment for varying levels of missing peaks (proportion of full data set). A higher F1 score means that the method is able to recover more of the original set of significant pathways correctly, even in the presence of missing peaks. The results show that PALS performs better than the alternatives on real data.

At the highest proportion of missing peaks (0.8) with the Plasma data, PALS achieves a mean *F*_1_ score of 0.48, while ORA and GSEA obtains 0.20 and 0.08, respectively (Supplementary Table S8). Similarly, on the CSF data, PALS achieves an F1 score of 0.60 while ORA and GSEA obtains 0.50 and 0.16, respectively. ORA on average returns higher mean precision values among the benchmarked methods, but the better *F*_1_ score of PALS can be attributed to its generally superior recall performance while still offering competitive precision in comparison to ORA. The results here suggest that even when there are many missing peaks, PALS is able to recover more of original pathways in the full data using a small fraction of peaks present in the original peak data, returning a higher number of true positives and fewer false positives and false negatives.

### 3.3 Analysis of User-defined Metabolite Sets

To demonstrate the analysis of user-defined metabolite sets, PALS was used to assess the activity levels of potentially unknown metabolites grouped into Molecular Families (MFs) through their fragmentation spectral similarities, and into Mass2Motifs based on shared fragment and neutral loss features.

For the MF analysis, we used the GNPS example data from the American Gut Project [21] where a subset was taken consisting of volunteers consuming differential amounts of plant-based food. 35 significantly changing MFs (p-value ≤ 0.05) containing 10 or more molecules were found between the case (eating ¿ 30 plant-based foods a week) and control (ļ 10 plant-based foods a week) groups. PALS prioritised one MF (p-value ≤ 0.001) of 23 molecules having steroid-related GNPS library hits that was found to be more abundant in the control group (for more details, see Supplementary Section S9). This could be due to the abundance of steroids in meat, fish, and eggs. Most other significant Molecular Families did not contain GNPS library hits but could represent potentially novel chemical classes.

For Mass2Motifs analysis, we used a dataset of 70 Rhamnacea species from two clades and various genera previously run through the GNPS-MS2LDA workflow [14]. In the original study, 25 Mass2Motifs were manually characterized and their distribution over the Rhamnaceae clades examined. Using PALS the statistical significance of the results can now be reported, while previous efforts relied on manual interpretation. Consistent with [14], PALS revealed that Mass2Motifs annotated with flavonoid-related substructures (i.e., rhamnocitrin, kaemfperol, flavonoid core framgent, and emodin) are all differently expressed between the Rhamnus and Ziziphus genera with features generally overrepresented in Rhamnus (for more details, see Supplementary Section S10). Similarly, the substructure set containing cyclopeptidic alkaloids was found to be overrepresented in Ziziphus. Differential metabolite sets for other genera could also be easily examined in PALS Viewer (for example the Xylose or Arabinose moiety substructure was found to be differentially expressed between Ventilago and Rhamnus). Finally, in the above-mentioned American Gut Project data, we could also detect 11 significant Mass2Motifs with one being related to ferulic acid (12 members) linked to plant-based foods (see Supplementary Section S9 for more details).

## 4 Discussion and Conclusion

In this work we describe PALS, a comprehensive system that can be used to prioritise metabolite sets, including but not limited to pathways, molecular families, and Mass2Motifs. It can be run in several modes: as a stand-alone tool, an interactive Web application or a library. From PALS, users obtain the ranking of changing pathways between experimental factors. Database queries that map compounds to pathways are integrated into the system using either KEGG or Reactome databases. User-defined metabolite sets can also be imported and analysed. A systematic comparison using synthetic and real data was performed against both ORA and GSEA, considered two of the most widely used methods for pathway ranking in metabolomics. The results show that the decomposition approach via PLAGE employed in PALS is consistently more robust in the presence of noisy peaks and missing peaks, a common problem in LC-MS peak data.

A weakness of all pathway methods is that it requires information on peak to formula annotation. This information has to be provided by the user, and it is assumed that the provided annotation is correct. This is not necessarily the case as metabolite identification remains an open problem in large-scale untargeted metabolomics [8, 7]. To our knowledge, this is a weakness of all pathway methods, except Mummichog [19] and PUMA [11] that combine annotation and activity prediction steps together. Future efforts on PALS will also aim to integrate a probabilistic model that assigns peaks to formulae based on mass accuracy and the prior connectivity of compounds in a pathway [24]. This is anticipated to improve the robustness of the method even further while allowing for uncertainties in formula annotations to be incorporated into pathway ranking.

## References

[1] Charras-Garrido, M. and Lezaud, P. (2013). Extreme value analysis: an introduction. Journal de la Société Française de Statistique, 154(2), 66–97.

[2] Chaudhary, K. and Roos, D. S. (2005). Protozoan genomics for drug discovery. Nature Biotechnology, 23(9), 1089–91.

[3] Chong, J. and Xia, J. (2018). MetaboAnalystR: an R package for flexible and reproducible analysis of metabolomics data. Bioinformatics, 34(24), 4313–4314.

[4] Chong, J. et al. (2018). Metaboanalyst 4.0: towards more transparent and integrative metabolomics analysis. Nucleic Acids Research, 46(W1), W486–W494.

[5] Creek, D. J. et al. (2015). Probing the metabolic network in bloodstream-form trypanosoma brucei using untargeted metabolomics with stable isotope labelled glucose. PLoS Pathogens, 11(3), e1004689.

[6] Croft, D. et al. (2013). The Reactome pathway knowledgebase. Nucleic Acids Research, 42(D1), D472–D477.

[7] da Silva, R. R. et al. (2015). Illuminating the dark matter in metabolomics. Proceedings of the National Academy of Sciences, 112(41), 12549–12550.

[8] Dunn, W. B. et al. (2013). Mass appeal: metabolite identification in mass spectrometry-focused untargeted metabolomics. Metabolomics, 9(1), 44–66.

[9] Ernst, M. et al. (2019). MolNetEnhancer: enhanced molecular networks by integrating metabolome mining and annotation tools. Metabolites, 9(7), 144.

[10] Gloaguen, Y. et al. (2017). PiMP my metabolome: an integrated, web-based tool for LC-MS metabolomics data. Bioinformatics, 33(24), 4007–4009.

[11] Hosseini, R. et al. (2020). Pathway-Activity Likelihood Analysis and Metabolite Annotation for Untargeted Metabolomics Using Probabilistic Modeling. Metabolites, 10(5), 183.

[12] Kamburov, A. et al. (2011). Integrated pathway-level analysis of transcriptomics and metabolomics data with IMPaLA. Bioinformatics, 27(20), 2917–2918.

[13] Kanehisa, M. and Goto, S. (2000). KEGG: Kyoto encyclopedia of genes and genomes. Nucleic Acids Research, 28(1), 27–30.

[14] Kang, K. B. et al. (2019). Comprehensive mass spectrometry-guided phenotyping of plant specialized metabolites reveals metabolic diversity in the cosmopolitan plant family rhamnaceae. The Plant Journal, 98(6), 1134–1144.

[15] Kessler, N. et al. (2013). MeltDB 2.0–advances of the metabolomics software system. Bioinformatics, 29(19), 2452–2459.

[16] Khatri, P. et al. (2012). Ten years of pathway analysis: current approaches and outstanding challenges. PLoS Computational Biology, 8(2), e1002375.

[17] Kind, T. and Fiehn, O. (2006). Metabolomic database annotations via query of elemental compositions: mass accuracy is insufficient even at less than 1 ppm. BMC Bioinformatics, 7(1), 234.

[18] Kluyver, T. et al. (2016). Jupyter notebooks-a publishing format for reproducible computational workflows. In Proceedings of the 20th International Conference on Electronic Publishing, pages 87–90.

[19] Li, S. et al. (2013). Predicting network activity from high throughput metabolomics. PLoS Computational Biology, 9(7), e1003123.

[20] Mantilla, B. S. et al. (2017). Proline metabolism is essential for trypanosoma brucei brucei survival in the tsetse vector. PLoS Pathogens, 13(1), e1006158.

[21] McDonald, D. et al. (2018). American gut: an open platform for citizen science microbiome research. MSystems, 3(3), e00031–18.

[22] Nothias, L. F. et al. (2019). Feature-based molecular networking in the GNPS analysis environment. bioRxiv, page 812404.

[23] Persicke, M. et al. (2012). MSEA: metabolite set enrichment analysis in the meltdb metabolomics software platform: metabolic profiling of corynebacterium glutamicum as an example. Metabolomics, 8(2), 310–322.

[24] Rogers, S. et al. (2008). Probabilistic assignment of formulas to mass peaks in metabolomics experiments. Bioinformatics, 25(4), 512–518.

[25] Subramanian, A. et al. (2005). Gene set enrichment analysis: a knowledge-based approach for interpreting genome-wide expression profiles. Proceedings of the National Academy of Sciences of the United States of America, 102(43), 15545–15550.

[26] Tarca, A. L. et al. (2013). A comparison of gene set analysis methods in terms of sensitivity, prioritization and specificity. PloS One, 8(11), e79217.

[27] Tomfohr, J. et al. (2005). Pathway level analysis of gene expression using singular value decomposition. BMC Bioinformatics, 6(1), 225.

[28] van Der Hooft, J. J. J. et al. (2016). Topic modeling for untargeted substructure exploration in metabolomics. Proceedings of the National Academy of Sciences, 113(48), 13738–13743.

[29] Vincent, I. M. et al. (2010). A molecular mechanism for eflornithine resistance in african trypanosomes. PLoS Pathogens, 6(11).

[30] Vincent, I. M. et al. (2016). Metabolomics identifies multiple candidate biomarkers to diagnose and stage human African trypanosomiasis. PLoS Neglected Tropical Diseases, 10(12), e0005140.

[31] Wang, M. et al. (2016). Sharing and community curation of mass spectrometry data with global natural products social molecular networking. Nature Biotechnology, 34(8), 828–837.

[32] Wishart, D. S. et al. (2012). HMDB 3.0—the human metabolome database in 2013. Nucleic Acids Research, 41(D1), D801–D807.

