## Supplementary Information for "Decomposing metabolite set activity levels with PALS"

[a]Glasgow Polyomics, University of Glasgow, Glasgow, G61 1BD, United Kingdom
[b]IBioIC, Strathclyde Institute of Pharmacy and Biomedical Sciences, University of Strathclyde, Glasgow, G1 1XQ, United Kingdom.
[c]Bioinformatics Group, Department of Plant Sciences, Wageningen University, Wageningen, The Netherlands.
[d]School of Computing Science, University of Glasgow, Glasgow G12 8RZ, United Kingdom.
[e]Centre for Synthetic and Systems Biology, School of Biological Sciences, University of Edinburgh, United Kingdom.


June 6, 2020

#### Supplementary Section S1: File format and data imputation

To use PALS, users have to provide information on peak intensities, peak annotations and the experimental design. Peak intensities should be provided as a matrix of individual peak intensities where the first column contains the peak IDs and further columns represent individual samples. When uploading a CSV file to the PALS Viewer or via the command line, the second line of this file should be used to indicate which groups this sample belongs to.

For example, the intensity matrix takes the form of:

```
peak_id,A001C.mzXML,A001P.mzXML,A002P.mzXML,A003C.mzXML,A004C.mzXML,A005C.mzXML,
    A008C.mzXML,A008P.mzXML,A009C.mzXML,A010C.mzXML,A010P.mzXML,A011C.mzXML
group, Control,Stage_2,Stage_2,Control,Control,Control,
    Control,Stage_1,Control,Control,Stage_1,Control
36186,612715072,408723328,356592704,575874816,627733440,399707168,
    621499008,545524352,522808352,557363328,541595328,476640800,531225312,535235328
36187,272679552,222055984,187961552,254896960,274777440,204597328,
    271147648,244269440,246383280,262514720,255993888,228511584
```

Data imputation is performed on the intensity matrix as follows: If all of the samples in a single experimental factor have intensities of zero these are replaced by the minimum intensity value (which can be set by the user); and if only some of the sample values in a factor are zero then these are replaced by the mean value of the non-zero samples in that factor. The data is subsequently transformed to log-2 base and standardised using the preprocessing module in Scipy [6] such that the intensity matrix has a zero mean and unit variance across the samples.

In addition to the peak intensities, users also provide a list of compound annotations assigned to peak features (peaks that do not have annotations will not be used for pathway analysis). As a result of the uncertainty in peak identification, multiple peak IDs may be mapped to multiple compound IDs and *vice versa*. As such, annotations are provided as another matrix having two columns. The first column (or DataFrame index) is the peak ID while the second column is the assigned metabolite annotation as either KEGG or ChEBI database IDs.

```
peak_id,entity_id
36883,C00111
37231,C00111
37309,C00661
37231,C19156
36368,C02718
37714,C05100
```

### Supplementary Section S2: Running PALS

PALS can be run in a variety of ways: from the command-line, from the Web interface (PALS Viewer) as well as imported directly as a Python library. Users should begin by first installing PALS using the following command: *pip install pals-pathway*, This retrieves the latest stable version of PALS from the Python Package Index.

#### S2.1 Running PALS from the command-line

To run PALS from the command-line, the script *pals/run.py* is used. This script accepts a number of parameters, documented here (* indicates required parameters):

```
usage: run.py [-h] --db {PiMP_KEGG,COMPOUND,ChEBI,UniProt,ENSEMBL}
              --comparisons COMPARISONS [COMPARISONS ...]
              [--min_replace MIN_REPLACE]
              [--species {Arabidopsis thaliana,Bos taurus,Caenorhabditis elegans,
              Canis lupus familiaris,Danio rerio,Dictyostelium discoideum,
              Drosophila melanogaster,Gallus gallus,Homo sapiens,Mus musculus,
              Oryza sativa,Rattus norvegicus,Saccharomyces cerevisiae,Sus scrofa}]
              [--use_all_reactome_pathways] [--connect_to_reactome_server]
              {PLAGE,ORA,GSEA} intensity_csv annotation_csv output_file
```

**method ***
: Pathway ranking method to use, e.g. PLAGE, ORA or GSEA.

**intensity_csv ***
: Input intensity CSV file (see Supplementary Section S1).

**annotation_csv ***
: Input annotation CSV file (see Supplementary Section S1).

**output_file ***
: Output pathway ranking file.

**–db ***
: The pathway database to use. Valid choices are as follows.

- *PiMP_KEGG*: KEGG compound database exported from PiMP.
- *COMPOUND*: Reactome compound database matching by KEGG ids.
- *ChEBI*: Reactome compound database matching by ChEBI ids.
- *UniProt*: Reactome protein database matching by UniProt ids.
- *ENSEMBL*: Reactome gene database matching by ENSEMBL ids.

Note that *PiMP_KEGG*, *COMPOUND* and *ChEBI* are for metabolomics use, while *UniProt* and *ENSEMBL* are for proteomics and transcriptomics use respectively and are not considered in this paper (refer to the project Web site for more information).

**–comparisons ***
: Specifies the comparisons to make, e.g. *–comparisons Stage_1/Control Stage_2/Control* to specify Stage 1 (case) vs control, as well as Stage_2 (case) vs control.

**–min_replace**
: The minimum intensity value for data imputation, e.g. *–min_replace 5000*. Defaults to 5000.

**–species**
: Species name for Reactome pathway query, e.g. *–species "Homo sapiens"*. Defaults to Homo Sapiens.

**–use_all_reactome_pathways**
: Whether to use all pathways for Reactome pathway query. If this option is not used, only metabolic pathways will be queried.

**–connect_to_reactome_server**
: Whether to connect to an instance of Neo4j server hosting Reactome database (online mode). If not specified, then offline mode (using a downloaded copy of selected Reactome pathways) will be used.

#### S2.2 Running PALS Viewer

PALS Viewer is a user-friendly graphical user interface to run PALS and analyse pathway ranking results as well as inspect significantly changing pathways. It can be run using the following command: *streamlit run pals/run_gui.py*. An online instance of PALS Viewer can also be accessed from the project Web site at `https://pals.glasgowcompbio.org/`. An example of PALS Viewer results for the CSF HAT data is shown below: **(A)** Activity results are shown in the Pathway Ranking table. Entries can be sorted and filtered by p-value threshold or the number of formula hits. **(B)** An example Reactome pathway selected from the Pathway Browser. Fold change values are mapped onto the pathway diagram using Reactome Analysis Service.

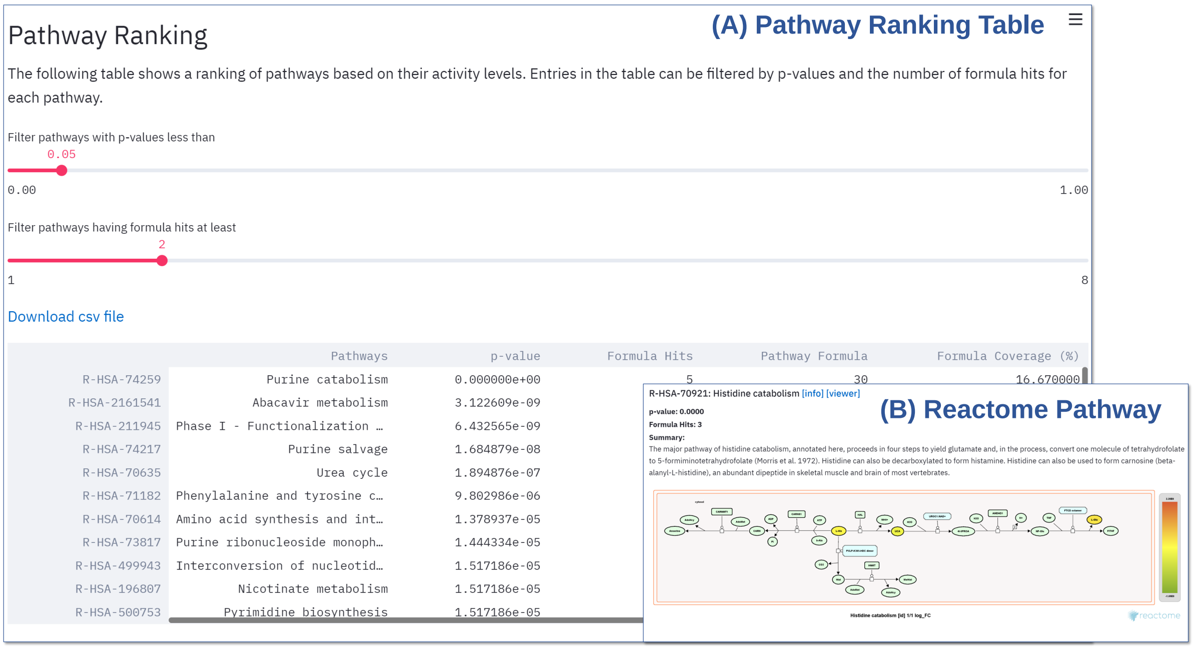

#### S2.3 Using PALS as a library

PALS can be imported as a Python library and incorporated into your own Python application. This is illustrated in the following code snippet (for additional documentation and tutorials, please refer to the project Web site):

```
from pals.PLAGE import PLAGE
from pals.ORA import ORA
from pals.GSEA import GSEA
from pals.common import *
from pals.feature_extraction import DataSource

# TODO: correctly initialise the following data structures for your data
# See Section S2.3.1 below.
int_df = pd.DataFrame()
annotation_df = pd.DataFrame()
experimental_design = {}

# Using Reactome pathways matching by KEGG ID
database_name = 'COMPOUND'

# If true, we limit to metabolic pathways only. Otherwise all pathways will be queried.
reactome_metabolic_pathway_only = True

# If true, we use online mode that queries Reactome on a local Neo4j server.
# Otherwise offline mode will be used (using downloaded database files).
reactome_query = True

# Minimum intensity value for data imputation
min_replace = 5000

ds = DataSource(int_df, annotation_df, experimental_design, database_name,
                reactome_species=reactome_species,
                reactome_metabolic_pathway_only=reactome_metabolic_pathway_only,
                reactome_query=reactome_query, min_replace=min_replace)

# choose a method
method = PLAGE(ds)
# method = ORA(ds)
# method = GSEA(ds)

df = method.get_pathway_df()
```

##### S2.3.1. Data structures

When PALS is used programatically, pandas DataFrames storing the intensity and annotation data, along with a dictionary describing the experimental design, can be passed directly to the program.

In the example above, *int_ df* is the intensity DataFrame containing peak intensity information described in Section S1 (with the second line of grouping information omitted). Similarly *annot_ df* is the annotation DataFrame containing peak annotations as described in Section S1. The experimental design data in *experimental_ design* contains information on ‘groups’, which relates all samples in a particular experimental factor together as well as ‘comparisons’, which describes the desired comparisons for the PALS analysis in terms of a case and a control. An example of this can be found below:

```
experimental_design = {
    'comparisons': [
        {'case': 'Stage_1', 'control': 'Control', 'name':  'Stage_1/Control'},
        {'case': 'Stage_2', 'control': 'Control', 'name': 'Stage_2/Control'},
        {'case': 'Stage_2', 'control': 'Stage_1', 'name': 'Stage_2/Stage_1'}
    ],
    'groups': {
        'Stage_1': [
            'A008P.mzXML',
            'A009P.mzXML',
            'A010P.mzXML'
        ],
        'Stage_2': [
            'A001P.mzXML',
            'A002P.mzXML',
            'A003P.mzXML'
        ],
        'Control': [
            'A001C.mzXML',
            'A002C.mzXML',
            'A003C.mzXML'
        ],
    }
}
```

#### Supplementary Section S3: Synthetic data generation

Synthetic pathway data is constructed to have differentially expressed intensity matrix (an example is shown in the figure below). Two groups (case and control) are included. The log intensity values of the control group is drawn from a normal (Gaussian) distribution with mean 20.0 and standard deviation 5.0, while for the case group, a normal distribution with mean 40.0 and standard deviation 5.0 is used. Each pathway is associated with the specified number of metabolites within the set: $2, 4, 6, 10, 20, 40, 80$. To simplify the problem of assigning peaks to metabolites, we assume a one-to-one correspondence between a peak and a metabolite (one metabolite produces exactly one peak). A synthetic pathway is labelled by the number of metabolites assigned to it (e.g. pathway *twenty* has 20 metabolites and therefore 20 peaks). In addition, 100 background pathways containing only noise (showing no significant changes between the case and control groups) were generated. The number of metabolites in a background pathway was randomly drawn with a uniform probability from 5 to 50, while the log intensity value in the random pathway is drawn from a normal distribution with mean 0 and standard deviation of 1. To simulate missing peaks, which often occurs in real data due to improper parameters used in peak picking or other preprocessing steps in the pipeline (e.g. setting an intensity filter threshold that is too low), peaks are also randomly removed from pathways with a uniform probability of 0.2. The total number of pathways evaluated in the synthetic data experiment is 107, composed of seven significantly changing pathways and 100 noisy pathways.

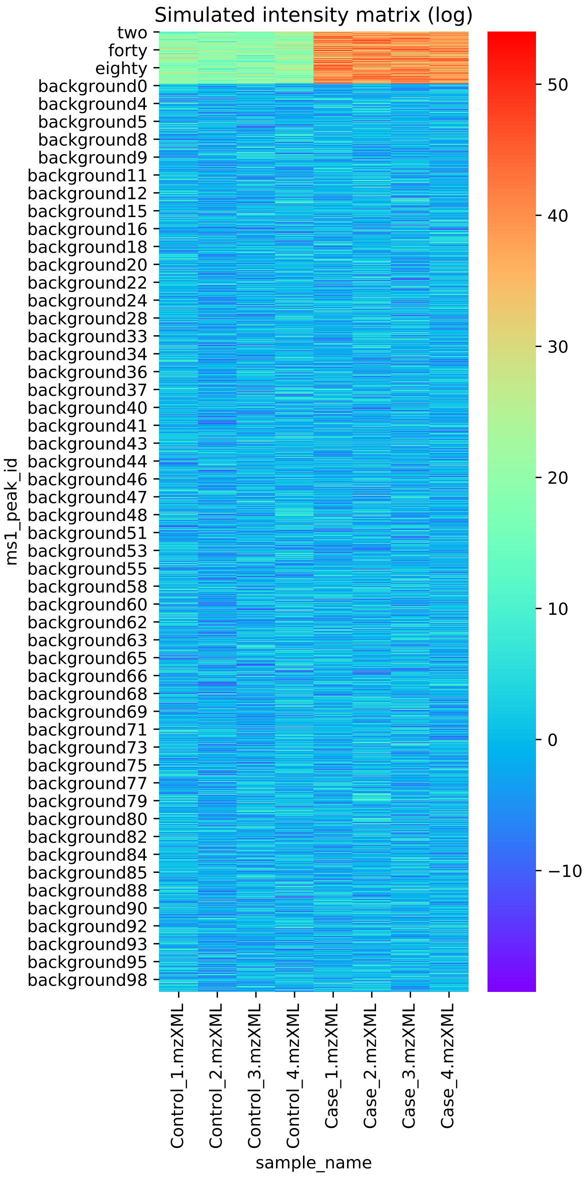

Simulated intensity matrix (log)
two
forty
eighty
background0
background4
background5
background8
background9
background11
background12
background15
background16
background18
background20
background22
background24
background28
background33
background34
background36
background37
background40
background41
background43
background44
background46
background47
background48
background51
background53
background55
background58
background60
background62
background63
background65
background66
background68
background69
background71
background73
background75
background77
background79
background80
background82
background84
background85
background88
background90
background92
background93
background95
background98
ms1_peak_id
Control_1.mzXML
Control_2.mzXML
Control_3.mzXML
Control_4.mzXML
Case_1.mzXML
Case_2.mzXML
Case_3.mzXML
Case_4.mzXML
sample_name
50
40
30
20
10
0
−10

#### Supplementary Section S4: Benchmark methods

The ORA method used for benchmarking is briefly summarised here: for each pathway, the number of significantly changing metabolites (above a p-value threshold of 0.05) is counted. Hypergeometric test is used to assess the probability of over-representation of significantly changing metabolites in that pathway. This procedure is repeated for all pathways, and the Benjamini-Hochberg correction is used to correct for multiple t-tests in the final results. For more details, refer to [2].

For GSEA, the GSEApy python package (`https://github.com/zqfang/GSEApy`), which implements the Gene-Set Enrichment Analysis algorithm in [5], was used. The steps in GSEA include the calculation of an enrichment score (ES). This is achieved by ranking metabolites according to the correlation of their peak intensity profiles to different experimental factors. Subsequently, a permutation test is performed to estimate the significance of the observed ES to the null hypothesis by randomly permuting factor labels, and correcting for multiple hypothesis testing by computing the false discovery rate. Following the original GSEA paper [5], the recommended number of 1000 permutations was used, as well as permuting the phenotype (sample) labels rather than the gene labels during permutation test. To produce the initial ranking of metabolites, we use the signal-to-noise ratio, which is also used by default in GSEA.

#### Supplementary Figure S5: Simulated Missing Peaks Results

In this section, the effect of introducing an increasing number of missing peaks from the pathways was introduced. For this experiment, the noise level was fixed to 100% in order to allow for an equal number of changing and random peaks. The fraction of peaks randomly missing from the data was 0.2, 0.4, 0.6 and 0.8, and the different pathway ranking methods were run 500 times for each dataset with missing peaks. The results in Figure 1 show that as as the number of missing peaks increases, PALS consistently returns lower mean p-values with smaller variances than ORA or GSEA. This suggests that PALS is generally more robust to missing peaks than the alternative methods tested.

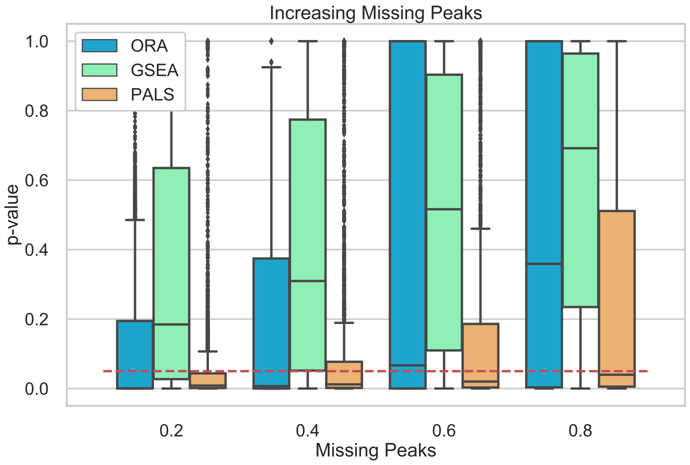


Figure 1: *Resulting p-values from removing peaks from significantly changing pathways. The fraction of peaks randomly removed from the pathways increases from 0.2 to 0.8 and ORA, GSEA and PALS are compared. The red dashed line indicates a p-value threshold of 0.05. The results show that PALS perform better returning lower p-values compared to the alternatives, even in the presence of a large number of missing peaks.*

These results were also supported by the $F_1$ score performance (Figure 2), where PALS generally performed best even when large number of peaks were missing from the data.

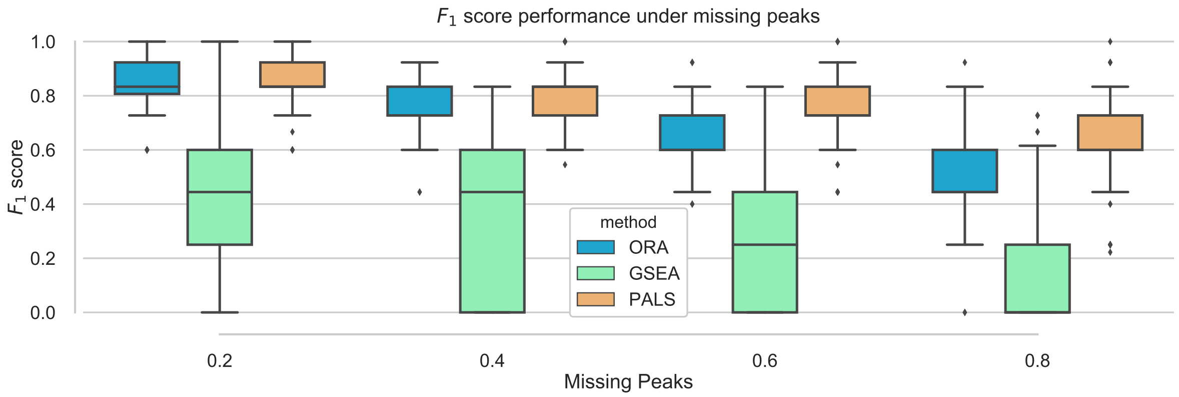


Figure 2: *Overall performance of the different pathway ranking methods with missing peaks. Distribution of the $F_1$ scores for each method under an increasing proportion of missing peaks (0.2 to 0.8) is shown. The results show PALS perform better returning greater $F_1$-scores at higher proportion of missing peaks.*

#### Supplementary Table S6: Top 30 ranking pathways from the HAT CSF dataset

| Pathway Name | AF | TPF | FC % | S1/S2 | S2/C | S1/C |
|---|---|---|---|---|---|---|
| Pantothenate and CoA biosynthesis | 11 | 25 | 44 | 0.0E+00 | 6.6E-26 | 9.7E-01 |
| Cyanoamino acid metabolism | 13 | 40 | 32.5 | 0.0E+00 | 2.8E-11 | 4.1E-02 |
| Arginine and proline metabolism | 23 | 79 | 29.11 | 0.0E+00 | 1.3E-09 | 6.8E-04 |
| Purine metabolism | 9 | 78 | 11.54 | 0.0E+00 | 2.4E-09 | 4.3E-01 |
| Alcoholism | 3 | 10 | 30 | 0.0E+00 | 2.6E-09 | 9.3E-02 |
| Aminoacyl-tRNA biosynthesis | 14 | 23 | 60.87 | 0.0E+00 | 5.5E-09 | 1.2E-03 |
| Cocaine addiction | 2 | 7 | 28.57 | 0.0E+00 | 5.8E-09 | 2.5E-01 |
| Amphetamine addiction | 2 | 9 | 22.22 | 0.0E+00 | 7.8E-09 | 2.6E-01 |
| Amyotrophic lateral sclerosis (ALS) | 2 | 10 | 20 | 0.0E+00 | 2.1E-08 | 2.6E-01 |
| Protein digestion and absorption | 14 | 42 | 33.33 | 0.0E+00 | 3.3E-08 | 3.1E-03 |
| Mineral absorption | 10 | 26 | 38.46 | 0.0E+00 | 2.0E-07 | 4.2E-02 |
| ABC transporters | 18 | 80 | 22.5 | 0.0E+00 | 2.3E-07 | 1.7E-02 |
| Anticonvulsants | 1 | 4 | 25 | 0.0E+00 | 2.8E-07 | 8.1E-01 |
| Alanine, aspartate and glutamate metabolism | 7 | 23 | 30.43 | 0.0E+00 | 4.1E-07 | 4.2E-01 |
| African trypanosomiasis | 1 | 7 | 14.29 | 0.0E+00 | 4.4E-07 | 8.3E-01 |
| Novobiocin biosynthesis | 2 | 25 | 8 | 0.0E+00 | 9.3E-07 | 7.5E-01 |
| Cysteine and methionine metabolism | 7 | 52 | 13.46 | 0.0E+00 | 1.3E-06 | 7.4E-01 |
| Neuroactive ligand-receptor interaction | 4 | 50 | 8 | 0.0E+00 | 1.5E-06 | 9.8E-01 |
| Indole alkaloid biosynthesis | 1 | 30 | 3.33 | 0.0E+00 | 2.0E-06 | 9.0E-01 |
| Histidine metabolism | 7 | 41 | 17.07 | 0.0E+00 | 5.2E-06 | 1.0E-03 |
| Phenylalanine, tyrosine and tryptophan biosynthesis | 9 | 30 | 30 | 0.0E+00 | 1.1E-05 | 5.4E-03 |
| Phenylalanine metabolism | 12 | 55 | 21.82 | 0.0E+00 | 1.9E-05 | 3.6E-02 |
| beta-Alanine metabolism | 7 | 31 | 22.58 | 0.0E+00 | 2.9E-05 | 2.4E-02 |
| Ubiquinone and other terpenoid-quinone biosynthesis | 6 | 56 | 10.71 | 0.0E+00 | 1.5E-04 | 1.1E-04 |
| Glutathione metabolism | 5 | 29 | 17.24 | 0.0E+00 | 4.3E-04 | 3.9E-02 |
| Caprolactam degradation | 7 | 19 | 36.84 | 2.5E-25 | 8.2E-01 | 1.6E-10 |
| Glycine, serine and threonine metabolism | 9 | 41 | 21.95 | 1.6E-22 | 8.4E-07 | 3.6E-01 |
| Bacterial chemotaxis | 1 | 5 | 20 | 1.5E-21 | 1.5E-05 | 9.9E-01 |
| Sphingolipid metabolism | 1 | 10 | 10 | 4.4E-21 | 2.4E-05 | 9.9E-01 |
| Glyoxylate and dicarboxylate metabolism | 8 | 48 | 16.67 | 4.8E-21 | 9.1E-06 | 7.1E-02 |

The top 30 best ranking pathways based on the PALS of the stage 1 (S1) compared to stage 2 (S2) in the cerebrospinal fluid (CSF) of patients with Human African Trypanosomiasis (HAT). The annotated formula (AF) found in the dataset and belonging to a particular pathway is shown along with the total formula expected in a pathway (TPF) and the percentage of the formula coverage (FC %). In the analysis, comparisons were made for between S1 and S2 along with S2 and S1 compared to the control (C) samples. The total number of KEGG pathways returned for this experiment was 162 and from these many of those involved in amino-acid metabolism were found to be highly significant.

#### Supplementary Table S7: Metabolites annotated in the KEGG aminoacyl-tRNA biosynthesis pathway

| Amino acid | Stage2/Stage1 (intensity) |
|---|---|
| L-Arginine | Significant decrease |
| L-Asparagine | Significant increase |
| L-Aspartate | Insignificant decrease |
| L-Glutamate | Significant increase |
| L-Glutamine | Insignificant decrease |
| L-Histidine | Significant decrease |
| L-Isoleucine | Insignificant decrease |
| L-Leucine | Significant decrease |
| L-Lysine | Significant decrease |
| L-Methionine | Insignificant decrease |
| L-Phenylalanine | Significant decrease |
| L-Proline | Significant increase |
| L-Serine | Significant decrease |
| L-Threonine | Significant decrease |
| L-Tryptophan | Significant decrease |
| L-Tyrosine | Significant decrease |
| L-Valine | Insignificant decrease |

The metabolites annotated in the KEGG aminoacyl-tRNA biosynthesis pathway in the CSF of HAT patients. Some metabolites show a significant increase or decrease, while others show an insignificant decrease in intensities between stage 1 and stage 2. All of the metabolite peaks were identified using in-house standards apart from L-Tyrosine that was identified through fragmentation and L-Aspartate for which no identification (only annotation was) was obtained.

#### Supplementary Table S8: Precision and recall on real HAT data

| Data | Missing Peaks | Method | Mean Prec. | Mean Recall | Mean $F_1$ |
|---|---|---|---|---|---|
| Plasma | 0.2 | ORA | 0.83 | 0.71 | 0.74 |
| | | GSEA | 0.71 | 0.22 | 0.30 |
| | | PALS | **0.86** | **0.88** | **0.87** |
| | 0.4 | ORA | **0.77** | 0.53 | 0.59 |
| | | GSEA | 0.55 | 0.18 | 0.24 |
| | | PALS | **0.77** | **0.76** | **0.76** |
| | 0.6 | ORA | **0.77** | 0.32 | 0.42 |
| | | GSEA | 0.43 | 0.12 | 0.16 |
| | | PALS | 0.69 | **0.61** | **0.64** |
| | 0.8 | ORA | 0.53 | 0.14 | 0.20 |
| | | GSEA | 0.20 | 0.06 | 0.08 |
| | | PALS | **0.61** | **0.42** | **0.48** |
| CSF | 0.2 | ORA | **0.95** | 0.87 | 0.91 |
| | | GSEA | 0.52 | 0.46 | 0.41 |
| | | PALS | 0.94 | **0.91** | **0.93** |
| | 0.4 | ORA | **0.92** | 0.75 | 0.82 |
| | | GSEA | 0.52 | 0.37 | 0.34 |
| | | PALS | 0.91 | **0.83** | **0.86** |
| | 0.6 | ORA | **0.90** | 0.60 | 0.71 |
| | | GSEA | 0.41 | 0.32 | 0.29 |
| | | PALS | 0.87 | **0.69** | **0.76** |
| | 0.8 | ORA | **0.87** | 0.38 | 0.50 |
| | | GSEA | 0.27 | 0.17 | 0.16 |
| | | PALS | 0.81 | **0.49** | **0.60** |

Mean precision, recall and $F_1$ score for the different methods under various missing peaks proportion on the Plasma and CSF data. The highest values (and ties) for precision, recall and $F_1$ score for each experimental setting is highlighted in bold.

### Supplementary Section S9: Analysis of Differentially Expressed Molecular Families and Mass2Motifs from the AGP Dataset

PALS was run on the following GNPS-FMBN [4] results from a previous analysis of a subset from the American Gut Project (AGP) dataset [3] comparing volunteers who eat differential amounts of plant-based food. The case group was selected to be those who eat more than 30 plant-based foods a week, while the control consists of those eating less than 10 plant-based foods a week.

For analysis, the GNPS-FBMN data available from `https://gnps.ucsd.edu/ProteoSAFe/status.jsp?task=0a8432b5891a48d7ad8459ba4a89969f` was used to extract MS1 peak table and grouping information. Sample metadata CSV is provided at `https://github.com/glasgowcompbio/PALS/raw/master/notebooks/test_data/AGP/AG_Plants_extremes_metadata_df.csv`. Analysis was performed using PALS Viewer at `https://pals.glasgowcompbio.org/app`. The following Jupyter notebook can also be used to perform the same analysis: `https://github.com/glasgowcompbio/PALS/blob/master/notebooks/GNPS_analysis.ipynb`.

In total, 35 significantly changing MFs containing 10 or more molecules were found to be DE between case and control groups. We found a notable Molecular Family containing steroid-related molecules of interest that is statistically significant (p-value $\leq 0.001$). This is plotted below, with the corresponding GNPS cluster id labelled green in the plot axes and also listed in the following table.

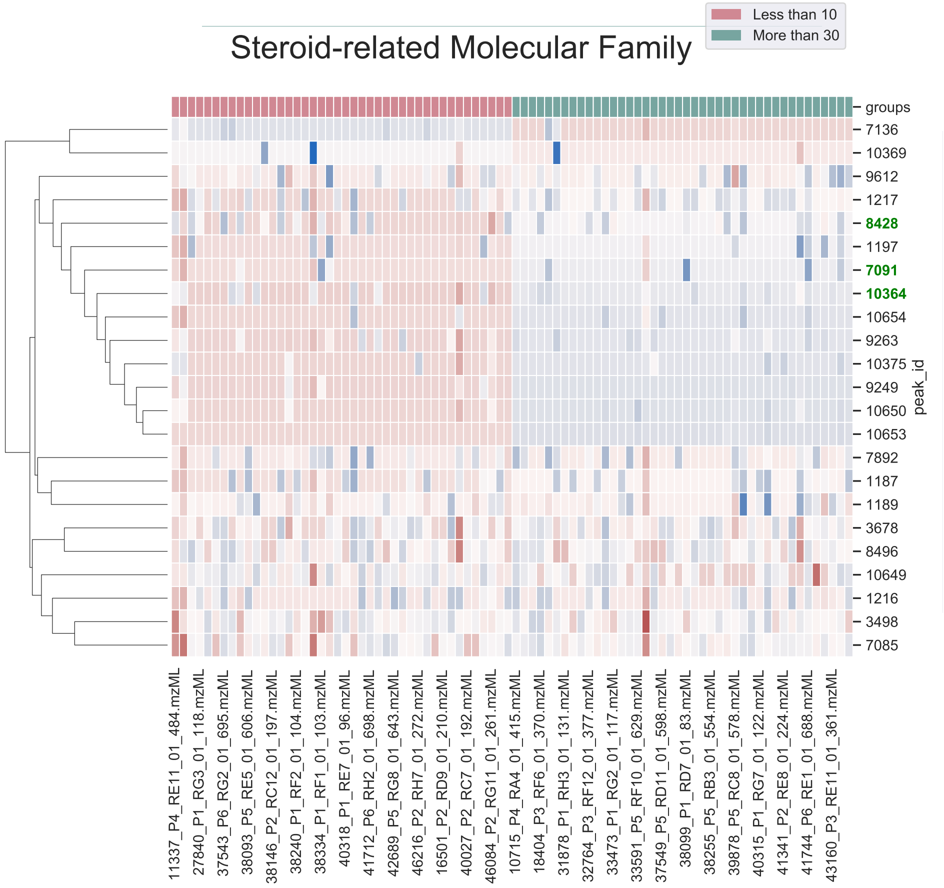

| id | LibraryID | m/z | RT | Intensity | no_spectra |
|---|---|---|---|---|---|
| 1187 | | 357.2056 | 3.0498 | 0.0377 | 61 |
| 1189 | | 385.2352 | 3.4066 | 0.0058 | 85 |
| 1197 | | 343.2264 | 3.6391 | 0.0185 | 28 |
| 1216 | | 367.2269 | 3.3133 | 0.0036 | 86 |
| 1217 | | 339.1964 | 3.0423 | 0.0038 | 109 |
| 3498 | | 769.4667 | 3.3526 | 0.001 | 174 |
| 3678 | | 399.3253 | 5.7595 | 0.0008 | 159 |
| 7085 | | 713.4052 | 3.0489 | 0.0031 | 176 |
| 7091 | Spectral Match to Mestranol from NIST14 | 311.2008 | 3.054 | 0.0012 | 24 |
| 7136 | | 343.2255 | 3.2039 | 0.0002 | 25 |
| 7892 | | 385.2367 | 3.1743 | 0.0133 | 68 |
| 8428 | adrenosterone | 301.1801 | 2.7956 | 0.0034 | 98 |
| 8496 | | 383.3314 | 8.3489 | 0.0076 | 196 |
| 9249 | | 343.2264 | 3.7812 | 0.0033 | 25 |
| 9263 | | 369.241 | 4.2581 | 0.0005 | 56 |
| 9612 | | 371.2578 | 4.3831 | 0.0082 | 90 |
| 10364 | Spectral Match to Boldione from NIST14 | 285.1853 | 3.1896 | 0.0009 | 52 |
| 10369 | | 413.3041 | 5.2675 | 0.0006 | 16 |
| 10375 | | 313.2165 | 3.7372 | 0.0004 | 28 |
| 10649 | | 685.4481 | 3.7336 | 0.0011 | 124 |
| 10650 | | 369.2419 | 4.1228 | 0.0019 | 30 |
| 10653 | | 341.2097 | 3.6374 | 0.0002 | 19 |
| 10654 | | 325.2157 | 3.6832 | 0.0004 | 17 |

‘ Finally for MS2LDA analysis, the AGP results for the FBMN workflow was further ran through the MS2LDA workflow on GNPS. This GNPS-MS2LDA result is available from `https://gnps.ucsd.edu/ProteoSAFe/status.jsp?task=7c34badae00e43bc87b195a706cf1f43`. The MS1 peak table from the original FBMN result was used for this analysis, as well as the provided metadata CSV. A significantly changing ferulic-acid related motif (p-value $\leq 0.001$) can be found below.

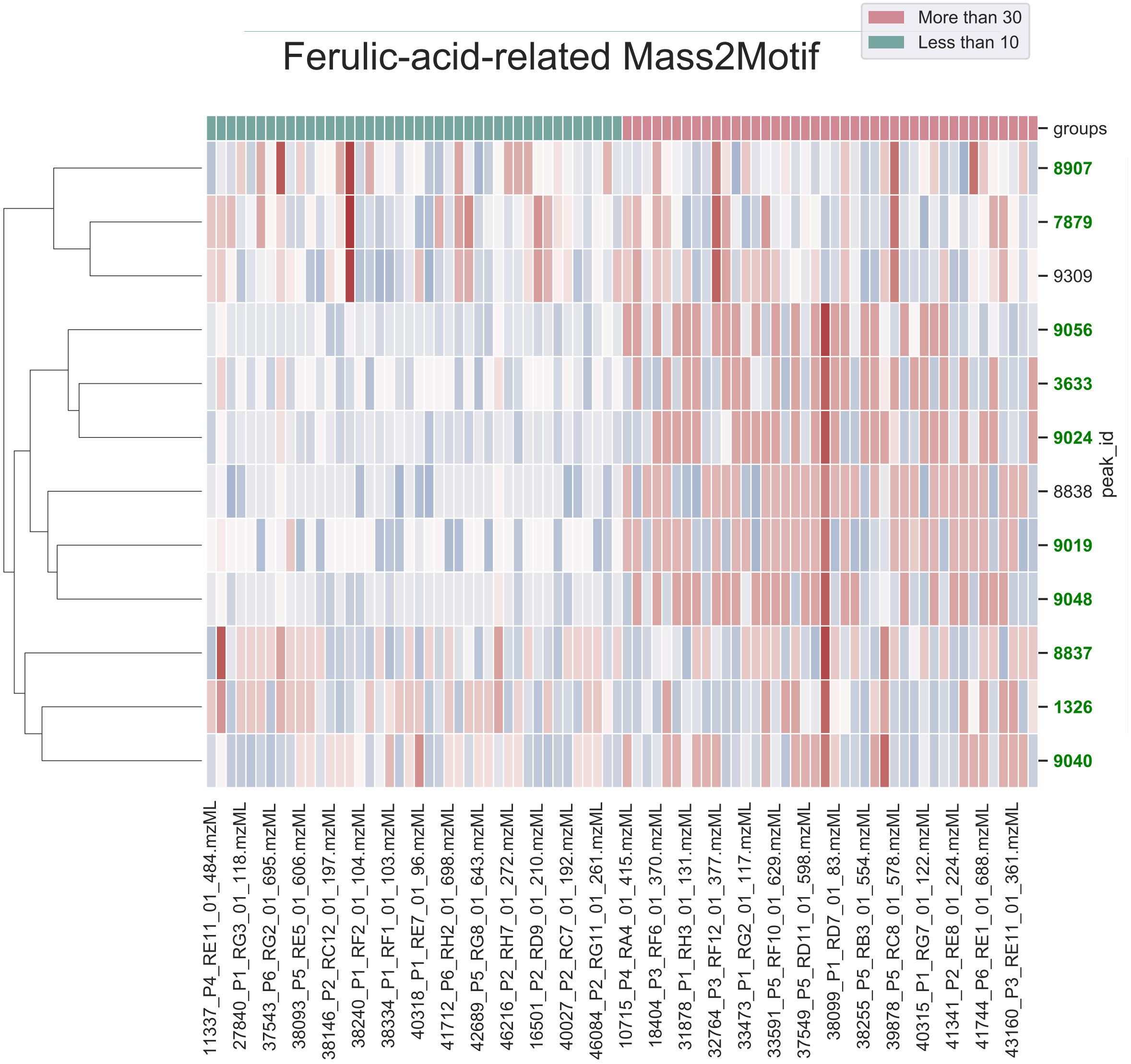

| id | LibraryID | m/z | RT | Intensity | no_spectra |
|---|---|---|---|---|---|
| 1326 | Spectral Match to Curcumin from NIST14 | 369.1335 | 4.0253 | 0.0322 | 145 |
| 3633 | Spectral Match to Curcumin from NIST14 | 369.1336 | 3.1547 | 0.0024 | 97 |
| 7879 | Spectral Match to 3-Hydroxy-4-methoxycinnamic acid from NIST14 | 195.0655 | 2.475 | 0.002 | 192 |
| 8837 | Spectral Match to 3-Hydroxy-4-methoxycinnamic acid from NIST14 | 177.0538 | 4.0199 | 0.0005 | 127 |
| 8838 | | 371.149 | 3.8366 | 0.0046 | 49 |
| 8907 | MoNA:3697220 Feruloyltyramine | 265.1549 | 0.889 | 0.0015 | 177 |
| 9019 | NCGC00095321-06!(1E,4Z,6E)-5-hydroxy-1,7-bis(4-hydroxy-3-methoxyphenyl)hepta-1,4,6-trien-3-one | 369.1332 | 4.275 | 0.0016 | 85 |
| 9024 | (1R,3R,4S,5R)-1,3,4-trihydroxy-5-[(E)-3-(4-hydroxyphenyl)prop-2-enoyl]oxycyclohexane-1-carboxylic acid | 339.123 | 3.147 | 0.0005 | 103 |
| 9040 | Spectral Match to Curcumin from NIST14 | 369.1331 | 3.877 | 0.0031 | 128 |
| 9048 | Spectral Match to Curcumin from NIST14 | 369.1331 | 4.4907 | 0.0007 | 84 |
| 9056 | NCGC00168971-02_C17H20O9_(1R,3R,4S,5R)-1,3,4-Trihydroxy-5-{[(2E)-3-(4-hydroxy-3-methoxyphenyl)-2-propenoyl]oxy}cyclohexanecarboxylic acid | 369.1335 | 3.3287 | 0.0003 | 105 |
| 9309 | | 177.0552 | 2.4861 | 0.0011 | 177 |

#### Supplementary Section S10: Analysis of Differentially Expressed Mass2Motifs from the Rhamnaceae Dataset

Using PALS Viewer, activity level analysis was run on the results of GNPS-MS2LDA workflow from [1] containing 25 Mass2Motifs that had been manually characterized and their distribution over the Rhamnaceae clades was examined.

The GNPS-MS2LDA data can be found from `https://gnps.ucsd.edu/ProteoSAFe/status.jsp?task=b33b2697e7924ee1920dba207ed57733`. To perform this analysis, users also need to upload the MS1 peak table (`https://github.com/glasgowcompbio/PALS/raw/master/notebooks/test_data/Rhamnaceae/171205_71extracts_MS1peaktable_MS2LDA_comma.csv`, as well as the metadata CSV describing which mzML files belong to which genera (`https://github.com/glasgowcompbio/PALS/raw/master/notebooks/test_data/Rhamnaceae/MetaData_Rhamnaceae.csv`).

We performed a comparison between the Rhamnus (control) and Ziziphus (case) genera. The results, shown in the following table, revealed that Mass2Motifs annotated with flavonoid-related substructures (i.e., rhamnocitrin, kaemfperol, flavonoid core framgent, and emodin) are all differently expressed between the Rhamnus and Ziziphus genera. The results here are consistent with the original study in [1],

| **Mass2Motif** | **p-value** | **No. of members** |
|---|---|---|
| rhamn_motif_130.m2m [Kaempferol] | 0.000000E+00 | 68 |
| rhamn_motif_121.m2m [CHOOH loss - indicative for underivatized carboxylic acid group] | 0.000000E+00 | 29 |
| rhamn_motif_140.m2m [Flavonoid core fragments (m/z 151)] | 0.000000E+00 | 25 |
| motif_81 | 0.000000E+00 | 24 |
| rhamn_motif_141.m2m [Emodin related Motif] | 0.000000E+00 | 21 |
| motif_50 | 0.000000E+00 | 12 |
| rhamn_motif_163.m2m [Glycosyl moiety] | 4.326859E-66 | 66 |
| rhamn_motif_40.m2m [Emodin related Motif] | 8.907510E-65 | 18 |
| rhamn_motif_164.m2m [rhamnocitrin-related] | 1.215099E-19 | 17 |
| motif_98 | 3.335502E-17 | 11 |
| rhamn_motif_172.m2m [CO2 loss] | 3.818233E-16 | 44 |
| motif_103 | 3.091902E-13 | 20 |
| motif_38 | 1.665557E-12 | 14 |
| motif_127 | 9.432444E-12 | 13 |
| motif_82 | 4.864967E-11 | 13 |
| rhamn_motif_28.m2m [coumaric acid-related] | 9.965255E-11 | 23 |
| rhamn_motif_87.m2m [(epi)ceanothic acid-related] | 1.021007E-09 | 27 |
| rhamn_motif_51.m2m [Cyclopeptide alkaloids] | 1.164000E-09 | 16 |
| rhamn_motif_167.m2m [Flavonoid core fragment (m/z 152)] | 1.577084E-09 | 33 |
| rhamn_motif_120.m2m [Coumaric acid - H2O] | 1.594465E-09 | 50 |
| motif_88 | 1.607455E-09 | 14 |
| rhamn_motif_179.m2m [Rhamnetin (=7-methylquercetin)] | 8.683922E-09 | 19 |
| motif_79 | 1.852689E-08 | 21 |
| rhamn_motif_153.m2m [CO2 loss] | 4.288013E-08 | 39 |
| rhamn_motif_169.m2m [coumaric acid related] | 5.868700E-08 | 11 |
| rhamn_motif_108.m2m [CHOOH loss - indicative for underivatized carboxylic acid group] | 1.169172E-07 | 15 |
| rhamn_motif_60.m2m [CO2/H2O loss] | 1.387670E-07 | 26 |
| motif_72 | 4.529178E-07 | 16 |
| rhamn_motif_165.m2m [ceanothic acid A-ring CO2 loss] | 1.318498E-06 | 24 |
| rhamn_motif_33.m2m [Xyl or Ara moiety] | 3.280194E-06 | 25 |
| motif_84 | 6.478872E-06 | 11 |
| motif_125 | 1.197932E-05 | 12 |
| motif_44 | 1.593861E-05 | 83 |
| rhamn_motif_48.m2m [Cyclopeptide alkaloids] | 2.034794E-05 | 12 |
| rhamn_motif_148.m2m [Cyclopeptide alkaloids] | 2.206457E-05 | 25 |
| motif_107 | 6.337228E-05 | 11 |
| rhamn_motif_117.m2m [protocatechuoyl-related] | 9.157846E-05 | 19 |
| motif_92 | 1.285183E-04 | 31 |
| motif_74 | 1.361637E-04 | 17 |
| rhamn_motif_191.m2m [vanilloyl-related] | 2.172679E-04 | 25 |
| motif_75 | 2.500638E-04 | 32 |
| rhamn_motif_34.m2m [Sugar (Glc) Loss] | 2.780931E-04 | 29 |
| motif_106 | 4.134637E-03 | 13 |
| motif_120 | 4.561325E-03 | 14 |
| rhamn_motif_64.m2m [Norrubrofusarin-related] | 5.617434E-03 | 13 |

### References

[1] Kang, K. B. *et al.* (2019). Comprehensive mass spectrometry-guided phenotyping of plant specialized metabolites reveals metabolic diversity in the cosmopolitan plant family rhamnaceae. *The Plant Journal*, **98**(6), 1134–1144.

[2] Khatri, P. *et al.* (2012). Ten years of pathway analysis: current approaches and outstanding challenges. *PLoS Computational Biology*, **8**(2), e1002375.

[3] McDonald, D. *et al.* (2018). American gut: an open platform for citizen science microbiome research. *MSystems*, **3**(3), e00031–18.

[4] Nothias, L. F. *et al.* (2019). Feature-based molecular networking in the GNPS analysis environment. *bioRxiv*, page 812404.

[5] Subramanian, A. *et al.* (2005). Gene set enrichment analysis: a knowledge-based approach for interpreting genome-wide expression profiles. *Proceedings of the National Academy of Sciences of the United States of America*, **102**(43), 15545–15550.

[6] Virtanen, P. *et al.* (2019). SciPy 1.0–Fundamental Algorithms for Scientific Computing in Python. *arXiv e-prints*, page arXiv:1907.10121.
